# Microbial Diversity and Differentiating Factors of Cocoa Fermentation Systems: Nutritional Supplementation as a Modulation Strategy Assessed by Metabarcoding

**DOI:** 10.64898/2026.04.16.718758

**Authors:** Carlos Eduardo Hernandez, Alejandra Mencia, Frank Solano-Campos, Adriana Arciniegas Leal

## Abstract

Cocoa fermentation is a spontaneous microbe-driven process in which yeasts, lactic acid bacteria (LAB), and acetic acid bacteria (AAB) generate the flavor precursors that determine the sensory quality of chocolate. Although the microbial ecology of cocoa fermentation has been increasingly studied through culture-independent methods, the effect of targeted nutritional interventions on community structure within geographically defined production territories has received limited attention. Here, we employed dual-marker metabarcoding (16S rRNA V4 and ITS1) with Illumina NovaSeq 6000 sequencing to characterize bacterial and fungal communities during spontaneous fermentation of Trinitario cocoa beans subjected to amino acid and zinc supplementation in the Limón province of Costa Rica. Fifteen samples were collected at 0, 24, and 48 h from control, amino acid-supplemented, and zinc-supplemented fermentations, each in duplicate. The bacterial community comprised 292 amplicon sequence variants (ASVs) representing 88 genera across 15 phyla; the fungal community comprised 1,117 ASVs representing 248 genera across 9 phyla. Firmicutes and Proteobacteria dominated the bacterial fraction, with a pronounced shift from *Tatumella*-dominated fresh pulp toward *Weissella*- and *Leuconostoc*-rich assemblages during fermentation. Amino acid supplementation reduced Firmicutes at 48 h while favoring *Acetobacter* proliferation; zinc supplementation promoted Mucoromycota and *Wickerhamomyces* while sustaining *Liquorilactobacillus* abundance. Beta diversity analyses (Aitchison distance, weighted and unweighted UniFrac) confirmed significant compositional differences between treatments (PERMANOVA, *p* ≤ 0.01), although alpha diversity indices did not differ between individual treatment pairs. Sparse Estimation of Correlations among Microbiomes (SECOM) revealed structured co-occurrence networks, including positive associations between *Gluconobacter* and *Acetobacter* and negative associations between *Tatumella* and several AAB genera. Predicted functional profiles (PICRUSt2) showed no significant pathway-level differences. Taken together, these results show that nutritional supplementation can reshape microbial community composition without reducing overall diversity. This provides a viable approach for steering fermentation outcomes in cocoa-producing territories that seek quality differentiation.

## 1. Introduction

Cocoa (*Theobroma cacao* L.) fermentation is a complex, spontaneous bioprocess in which the coordinated metabolic activities of yeasts, lactic acid bacteria (LAB), and acetic acid bacteria (AAB) transform the sugar-rich pulp surrounding the beans into organic acids, alcohols, esters, and aldehydes that serve as precursors of chocolate flavor (De Vuyst and Weckx, 2016; Schwan et al., 2014). The characteristic microbial succession, from an initial yeast-dominated anaerobic phase through LAB acidification to aerobic AAB oxidation, has been documented in major cocoa-producing regions. Yet considerable variability in community composition and metabolic output persists between farms, seasons, and geographical origins (Papalexandratou et al., 2011; Camu et al., 2007; Lefeber et al., 2011).

This variability has given rise to the concept of “microbial terroir,” analogous to the well-established notion in viticulture whereby indigenous microbial populations contribute region-specific sensory attributes to the final product (Bokulich et al., 2014). Recent metagenomic and metabarcoding studies have shown that cocoa fermentation microbiomes exhibit biogeographic patterns: certain yeast species such as *Hanseniaspora guilliermondii* predominate in West Africa, whereas *Pichia kudriavzevii* and *Saccharomyces cerevisiae* are more prevalent in Latin American fermentations (Campos et al., 2025; Díaz-Muñoz et al., 2021). A meta-analysis of 60 studies identified over 1,700 microbial species across Brazilian, Ivorian, and Ghanaian fermentations, but was unable to establish robust correlations between community composition and geographical origin, highlighting the stochastic nature of spontaneous fermentation and the need for controlled interventions (Taylor et al., 2022). Ouattara and Niamké (2021) mapped the functional diversity of fermentative microbiota across twelve cocoa-producing regions in Côte d’Ivoire and found that while acid-producing bacteria were relatively conserved, yeast populations varied considerably between regions, suggesting that the fungal community may be a primary driver of terroir-linked flavor differentiation.

Delimited cocoa-producing territories are regions with defined agro-ecological characteristics that contribute to the distinctive quality of their products. Understanding and harnessing microbial diversity during fermentation in these territories represents an opportunity for quality differentiation and value addition. Geographical indications (GIs) and denomination-of-origin frameworks for cocoa increasingly require evidence linking production practices to sensory quality (Hernandez and Granados, 2021). However, the factors that shape microbial community structure within specific territories remain insufficiently characterized, particularly with regard to manageable production variables.

Nutritional supplementation during fermentation represents one such controllable intervention. Zinc is an essential cofactor for over 300 enzymes in *S. cerevisiae*, including alcohol dehydrogenases critical to ethanol production, and its availability directly influences yeast vitality, stress tolerance, and the biosynthesis of higher alcohols and esters via the Ehrlich pathway (De Nicola et al., 2007; Zhao and Bai, 2012; Walker, 2004). Under zinc limitation, genes involved in branched-chain amino acid biosynthesis (*ILV2, ILV3, BAT2*) are down-regulated, reducing the flux toward desirable flavor compounds such as isoamyl acetate (De Nicola et al., 2007). Amino acid supplementation, in turn, enhances volatile aroma compound production through the Ehrlich pathway. Amino acids such as leucine, phenylalanine, and valine are transaminated, decarboxylated, and reduced to form higher alcohols. These are then esterified to produce fruity and floral esters (Hazelwood et al., 2008; Procopio et al., 2011).

Despite this established biochemical rationale, the effect of amino acid and zinc supplementation on the complex, multi-kingdom microbial communities of cocoa fermentation has not been systematically investigated using high-resolution culture-independent methods. Previous work in brewing and wine fermentation has demonstrated that mineral and nitrogen supplementation can alter microbial community dynamics and volatile profiles (Tenge and Geiger, 2001; Bell and Henschke, 2005), but the translatability of these findings to the ecologically distinct environment of cocoa fermentation is unknown.

In this study, we employed dual-marker amplicon sequencing (16S rRNA V4 for bacteria and archaea, ITS1 for fungi) to characterize the microbial diversity and community structure during spontaneous fermentation of Trinitario cocoa beans from a defined production territory in the Limón province of Costa Rica, under three conditions: unsupplemented control, amino acid supplementation, and zinc supplementation. Our objectives were to: (i) characterize the baseline microbial diversity and temporal dynamics of the fermentation microbiome in this territory; (ii) evaluate the effect of nutritional supplementation on bacterial and fungal community composition; (iii) identify taxa and inter-taxon correlations that respond differentially to supplementation; and (iv) assess predicted functional consequences of observed community shifts.

## 2. Materials and Methods

### 2.1. Study design and sample collection

The study evaluated the effect of amino acid and zinc supplementation on microbial diversity during fermentation of Trinitario cocoa beans in the Limón province of Costa Rica, a recognized fine-flavor cocoa territory. Freshly harvested cocoa pods were opened and the pulp-bean mass was placed in wooden fermentation boxes following standard farm practices. Three treatments were established: (i) unsupplemented control, (ii) amino acid supplementation, and (iii) zinc supplementation. Samples (∼20 g, ∼5 cocoa seeds with mucilage) were collected at t0 (fresh pulp), t24 (24 h), and t48 (48 h), in duplicate for each treatment, yielding 15 samples (3 fresh pulp replicates + 2 replicates × 2 time points × 3 conditions).

### 2.2. DNA extraction and quality control

DNA was isolated from cocoa bean samples following the protocol of Van de Voorde et al. (2023), with emphasis on removing contaminant plant material. After obtaining a cell pellet, total genomic DNA was extracted using the DNeasy PowerLyzer PowerSoil Kit (Qiagen, Hilden, Germany) according to the manufacturer’s instructions. DNA quality was confirmed by PCR amplification of the V3–V4 region of the 16S rRNA gene using primers 342F (5′-CTACGGGGGGCAGCAG-3′) and 806R (5′-GGACTACCGGGGTATCT-3′). Extracted DNA was shipped to Novogene Inc. (Sacramento, CA, USA) for library preparation and sequencing.

### 2.3. Amplicon library preparation and sequencing

Libraries targeting the V4 region of the 16S rRNA gene were prepared using primers 515F (5′-GTGCCAGCMGCCGCGGTAA-3′) and 806R (5′-GGACTACHVGGGTWTCTAAT-3′) (Caporaso et al., 2012). Libraries targeting the ITS1 region were prepared using primers ITS5-1737F (5′-GGAAGTAAAAGTCGTAACAAGG-3′) and ITS2-2043R (5′-GCTGCGTTCTTCATCGATGC-3′) (Bellemain et al., 2010). Both libraries were sequenced on the Illumina NovaSeq 6000 platform with 2 × 250 bp paired-end reads.

### 2.4. Bioinformatic processing

Primer sequences were removed using Cutadapt v3.5 (Martin, 2011). Amplicon sequence variants (ASVs) were inferred using DADA2 (Callahan et al., 2016) in R v4.3.2 (R Core Team, 2024). Taxonomic classification was performed with a naïve Bayesian classifier (Wang et al., 2007) using SILVA 138.1 (Quast et al., 2013) for bacteria and UNITE v10 (Abarenkov et al., 2024) for fungi. For bacteria, ASVs assigned to chloroplast, mitochondria, or eukaryotic lineages were removed; for fungi, ASVs not assigned to Fungi were removed. A reference phylogenetic tree was constructed by placing filtered bacterial ASVs into a 20,000-sequence reference alignment from the IMG database (Markowitz et al., 2012). This was accomplished using HMMER (Eddy, 2011), EPA-ng (Barbera et al., 2019), and GAPPA (Czech and Stamatakis, 2019), as implemented in PICRUSt2 (Douglas et al., 2020). Bacterial ASVs with poor phylogenetic placement were excluded.

### 2.5. Taxonomic profiling and differential abundance

Heatmaps of relative abundance at phylum, class, and genus levels were generated using ampvis2 (Andersen et al., 2018). Differential abundance testing was performed using ALDEx2 (Fernandes et al., 2014). Sparse Estimation of Correlations among Microbiomes (SECOM) was used to estimate inter-taxon linear correlations using the Pearson2 approach (Lin et al., 2022).

### 2.6. Alpha diversity analysis

Count tables were normalized by scaling with ranked subsampling (SRS; Beule and Karlovsky, 2020). Shannon, Chao1, Simpson, and Faith’s phylogenetic diversity (PD) indices were calculated using phyloseq v1.42.0 (McMurdie and Holmes, 2013) and picante v1.8 (Kembel et al., 2010). As the Shapiro–Wilk test indicated non-normal distributions, differences between treatments were evaluated by Kruskal–Wallis tests with post-hoc Dunn’s tests and Benjamini– Hochberg FDR correction.

### 2.7. Beta diversity analysis

Abundance and prevalence filters (≥4 reads in ≥10% of samples) were applied (Dhariwal et al., 2017; Motiei et al., 2020). Aitchison distances (Euclidean distance of centered log-ratio transformed data) were visualized by PCA (Gloor et al., 2017). Weighted and unweighted UniFrac distances were ordinated by PCoA using phyloseq. Community differences were assessed by ANOSIM, betadisper, and PERMANOVA using vegan v2.6-4 (Oksanen et al., 2010), with pairwise comparisons via RVAideMemoire v0.9 (Hervé, 2023).

### 2.8. Functional prediction

Metagenomic functions were predicted from bacterial ASVs using PICRUSt2 v2.5.2 (Douglas et al., 2020). KEGG Orthology abundances were tested for pathway-level differences between treatments using ggpicrust2 v1.7.3 (Yang et al., 2023) with DESeq2 (Love et al., 2014).

## 3. Results

### 3.1. Sequencing output and ASV recovery

The bacterial dataset contained 2,896,636 raw reads (mean 193,109 per sample). After quality filtering, merging, and chimera removal through DADA2, 2,721,844 reads (mean 181,456 per sample) were retained across 313 ASVs. Following taxonomic and phylogenetic filtering, 292 ASVs remained, representing 88 genera from 24 classes across 15 phyla. Fresh pulp harbored 109 ASVs, the control 182, the amino acid treatment 178, and the zinc treatment 163.

The fungal dataset contained 1,800,724 raw reads (mean 120,048 per sample), yielding 1,429,228 reads after processing (mean 95,282 per sample) across 1,695 ASVs. After filtering, 1,117 ASVs remained, representing 248 genera from 30 classes across 9 phyla. Fresh pulp harbored 881 ASVs, the control 160, the amino acid treatment 168, and the zinc treatment 137.

### 3.2. Bacterial taxonomic profiling

After abundance and prevalence filtering, the bacterial community comprised 3 phyla, 4 classes, and 28 genera (Figure 1). Proteobacteria (43 ASVs), Firmicutes (34 ASVs), and Actinobacteriota (5 ASVs) accounted for virtually all classified reads. Firmicutes was the most abundant phylum overall, followed by Proteobacteria, whereas Actinobacteriota remained at extremely low relative abundance (0.003–0.67%). The fresh pulp was dominated almost exclusively by Proteobacteria (99.26%). At the class level, Bacilli, Gammaproteobacteria, and Alphaproteobacteria predominated. The most abundant genera were *Weissella, Leuconostoc, Tatumella*, and *Acetobacter*.

**Figure 1.**
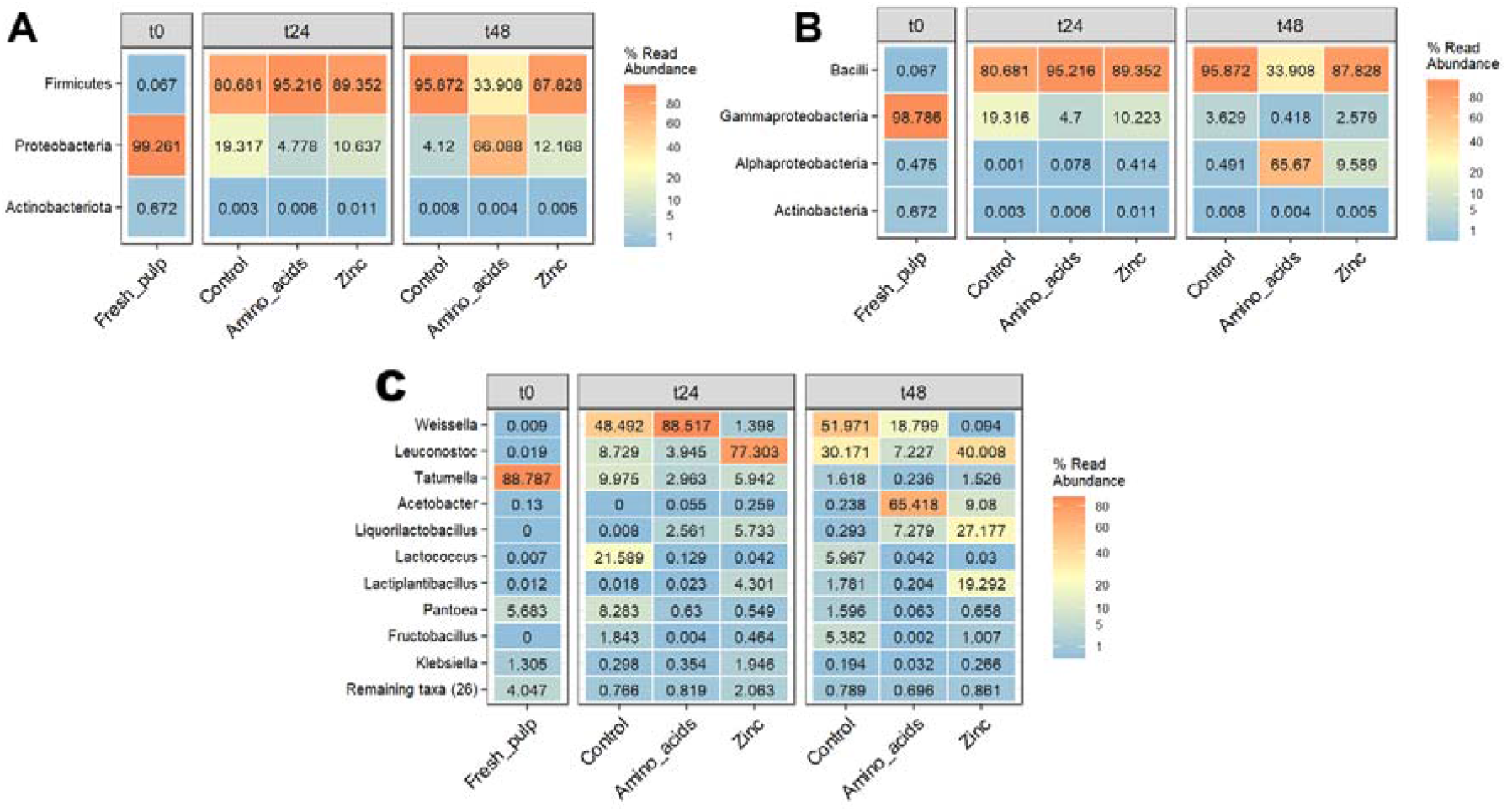
Heatmaps of the relative abundance of the three phyla (A), four classes (B), and top 10 genera (C) in the bacterial community of the cacao samples across treatments and time points. Color intensity reflects the percentage of read abundance.

### 3.3. Fungal taxonomic profiling

The filtered fungal community comprised 4 phyla, 14 classes, and 59 genera (Figure 2). Ascomycota (141 ASVs) and Basidiomycota (13 ASVs) dominated the community. Ascomycota was the most abundant phylum across all samples, while Basidiomycota reached 2.55% in fresh pulp. The dominant classes were Saccharomycetes, Sordariomycetes, Eurotiomycetes, and Leotiomycetes. The most abundant genera were *Hanseniaspora, Thielaviopsis, Pichia*, and *Cyphellophora*.

**Figure 2.**
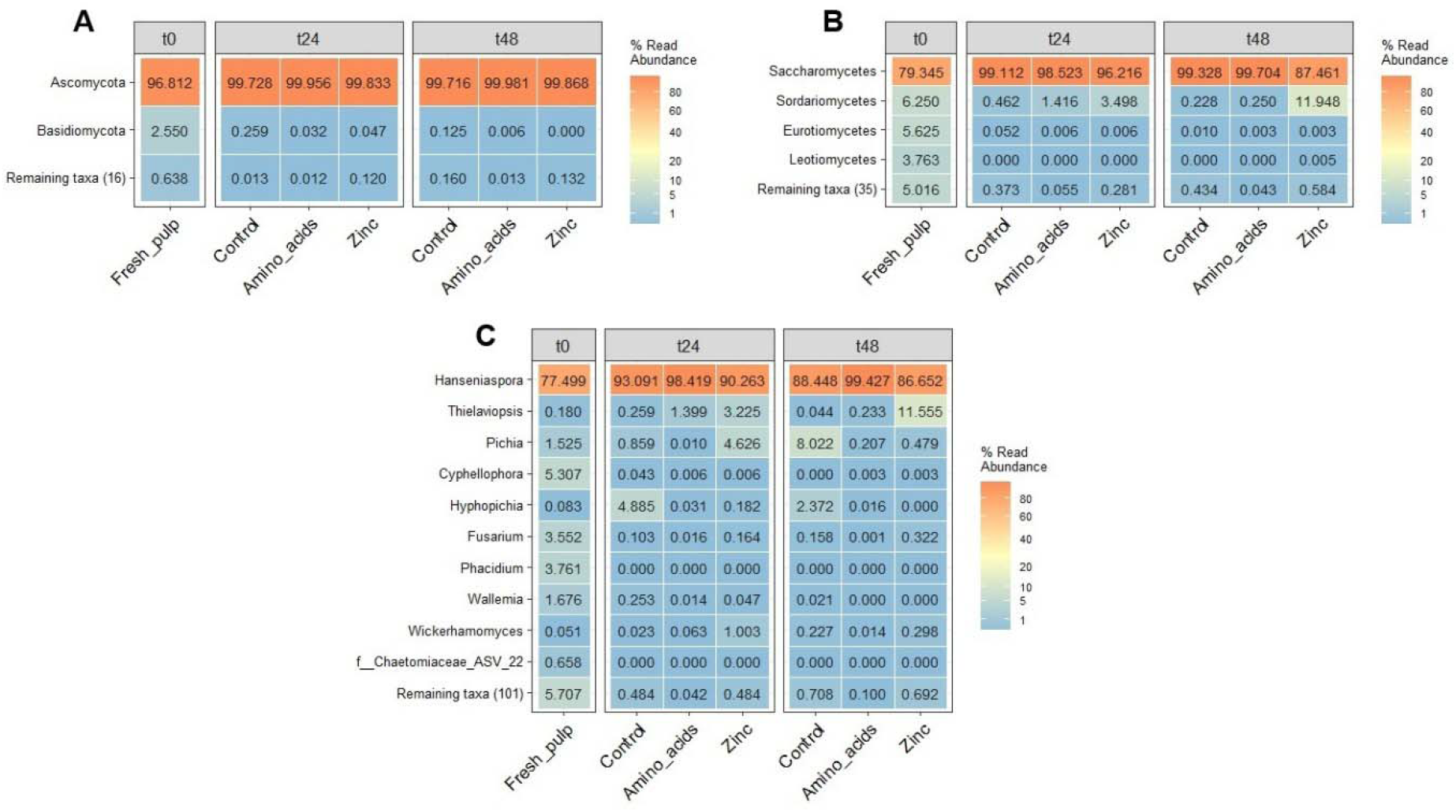
Heatmaps of the relative abundance of the two phyla (A), four classes (B), and top 10 genera (C) in the fungal community of the cacao samples across treatments and time points. Color intensity reflects the percentage of read abundance.

### 3.4. Differential abundance analysis

#### Bacterial community

ALDEx2 analysis identified significant differential abundance of Firmicutes and Proteobacteria between treatments. Firmicutes showed a significant reduction at 48 h under amino acid supplementation, with a concomitant increase in Proteobacteria. At the class level, Bacilli decreased and Alphaproteobacteria increased at 48 h in the amino acid treatment, while Gammaproteobacteria was highest in the control at 24 h. Of the 28 detected genera, 24 exhibited differential abundance. *Weissella* peaked in the amino acid treatment at 24 h; *Leuconostoc* peaked in the zinc treatment at 24 h; and *Tatumella* was highest in the control at 24 h. *Acetobacter* was markedly enriched in the amino acid treatment at 48 h (65.42% relative abundance), *Liquorilactobacillus* in the zinc treatment at 48 h, and *Lactococcus* in the control at 24 h. *Pantoea* and *Fructobacillus* were enriched in the control at 24 h and 48 h, respectively, while *Klebsiella* and *Lactiplantibacillus* were enriched in the zinc treatment at 24 h and 48 h.

#### Fungal community

All three fungal phyla showed significant differences. Basidiomycota was enriched in fresh pulp relative to later time points, while Ascomycota showed the opposite trend. Mucoromycota was increased in the zinc treatment at both 24 h and 48 h. At the class level, Eurotiomycetes was enriched in fresh pulp, Saccharomycetes increased during fermentation, and Mucoromycetes was elevated under zinc supplementation. Among 59 genera, 12 showed significant differential abundance, including 7 of the 10 most abundant: *Hanseniaspora* (lower in fresh pulp), *Cyphellophora, Fusarium*, and *Wallemia* (higher in fresh pulp), *Pichia* (lower in amino acid treatment), *Wickerhamomyces* (highest in zinc treatment at 24 h), and *Hyphopichia* (higher in control at 24 and 48 h).

### 3.5. Inter-taxon correlations

#### Bacteria

SECOM analysis revealed a significant negative correlation between Firmicutes and Actinobacteriota (Pearson2 = −0.86). Alphaproteobacteria and Gammaproteobacteria were strongly negatively correlated (−0.93), as were Bacilli and Actinobacteria (−0.86). Among genera, positive correlations were found between *Gluconobacter* and *Acetobacter* (0.84), *Leuconostoc* and *Lactiplantibacillus* (0.73), and *Weissella* and *Lactococcus* (0.78). Negative correlations were detected between *Tatumella* and *Acetobacter* (−0.82), *Gluconobacter* (−0.81), *Lactiplantibacillus* (−0.74), and *Leuconostoc* (−0.78). *Liquorilactobacillus* was negatively correlated with *Pantoea* (−0.86) and *Lactococcus* (−0.92).

#### Fungi

Ascomycota and Mucoromycota were negatively correlated (−0.87). Eurotiomycetes and Dothideomycetes showed a positive correlation (0.93), while Saccharomycetes and Mucoromycetes were negatively correlated (−0.89). At genus level, *Hanseniaspora* and *Wickerhamomyces* were negatively correlated (−0.79).

### 3.6. Alpha diversity

After normalization, 153 bacterial ASVs and 1,115 fungal ASVs were retained. Bacterial alpha diversity indices are shown in Table 1 and fungal indices in Table 2.

**Table 1.** Observed bacterial species richness and alpha diversity indices (mean ± SD) across treatments after normalization by scaling with ranked subsampling.

| Treatment | Observed | Chao1 | ACE | Shannon | Simpson | InvSimpson | Fisher | PD |
| --- | --- | --- | --- | --- | --- | --- | --- | --- |
| Fresh pulp | 62.67±12.9 | 89.14±23.95 | 96.36±26.11 | 0.93±0.26 | 0.38±0.11 | 1.64±0.31 | 8.93±2.15 | 26.26±3.94 |
| Control t24 | 40.50±2.12 | 54.44±14.94 | 49.40±5.60 | 1.53±0.36 | 0.66±0.17 | 3.37±1.66 | 5.37±0.32 | 11.34±0.48 |
| Control t48 | 59.00±5.66 | 82.87±15.17 | 75.94±7.83 | 1.47±0.29 | 0.62±0.12 | 2.77±0.86 | 8.31±0.93 | 15.16±1.44 |
| Amino t24 | 40.00±1.41 | 47.08±1.53 | 50.78±0.17 | 0.63±0.08 | 0.22±0.04 | 1.29±0.07 | 5.29±0.22 | 14.67±0.35 |
| Amino t48 | 49.50±0.71 | 61.77±3.31 | 67.60±7.48 | 1.86±0.02 | 0.79±0.01 | 4.75±0.17 | 6.77±0.11 | 13.93±0.91 |
| Zinc t24 | 52.50±10.61 | 95.12±32.00 | 74.50±26.62 | 1.11±0.13 | 0.40±0.04 | 1.68±0.13 | 7.26±1.70 | 16.61±3.78 |
| Zinc t48 | 56.00±1.41 | 62.38±4.07 | 64.41±3.84 | 1.82±0.09 | 0.75±0.02 | 4.01±0.33 | 7.81±0.23 | 14.23±0.80 |

**Table 2.**
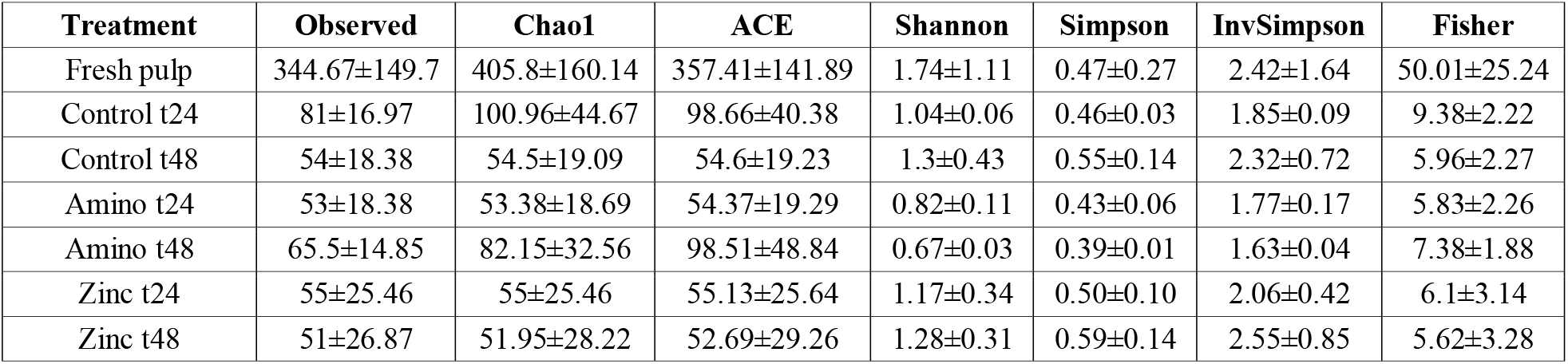
Observed fungal species richness and alpha diversity indices (mean ± SD) across treatments after normalization.

For bacteria, Kruskal–Wallis tests detected significant overall differences among treatments in Shannon (*p* = 0.047), Simpson (*p* = 0.044), and InvSimpson (*p* = 0.044), but not in Observed richness, Chao1, ACE, Fisher’s alpha, or PD (*p* > 0.05). Post-hoc Dunn’s tests did not identify significant pairwise differences between any two treatments (Figure 3). For fungi, no significant differences in any index were detected (*p* > 0.05).

**Figure 3.**
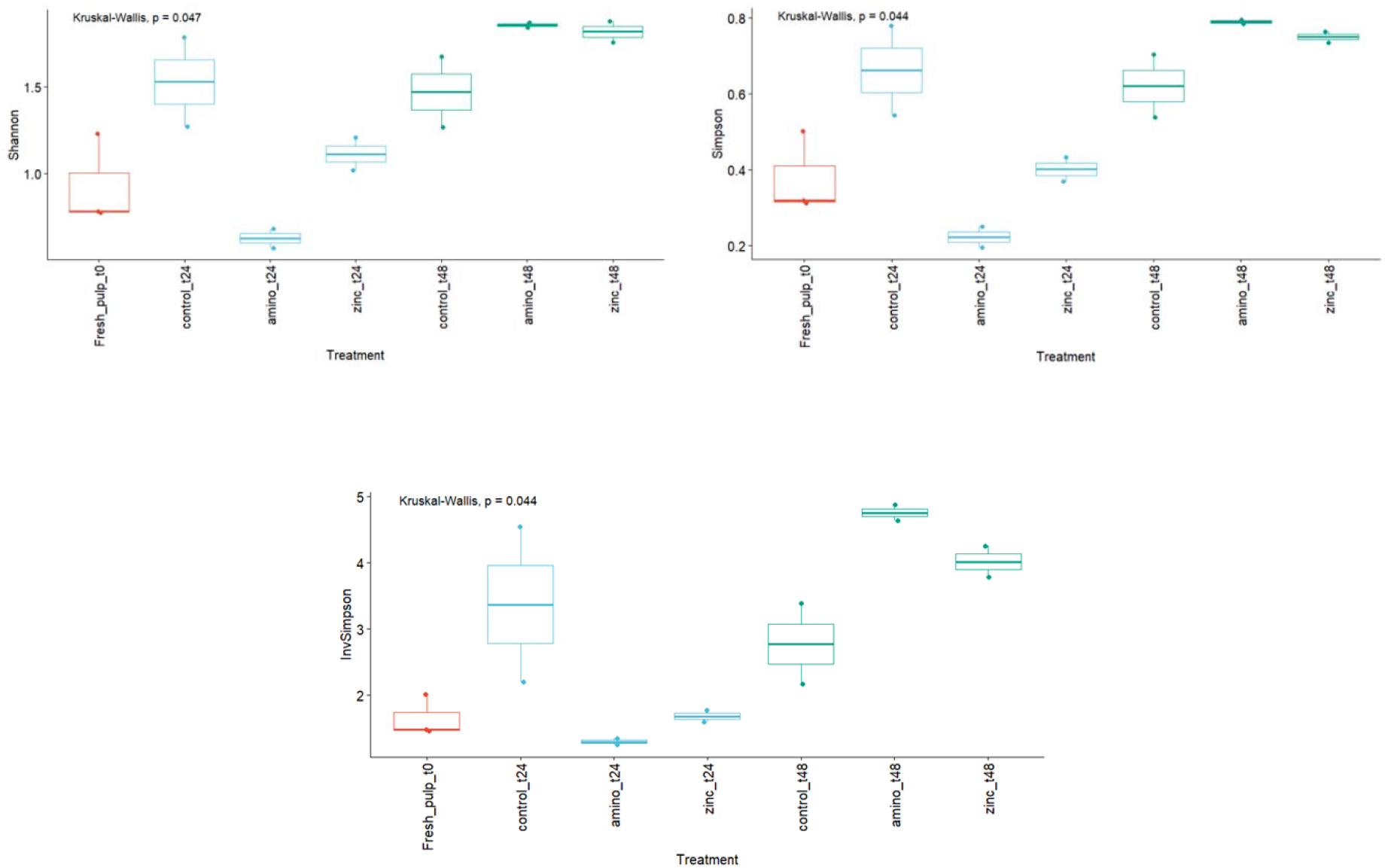
Bacterial alpha diversity represented as Shannon, Simpson, and InvSimpson indices across treatments. Count data were normalized by scaling with ranked subsampling. Significant overall differences were detected by Kruskal–Wallis test, but no pairwise differences were identified by post-hoc Dunn’s test.

### 3.7. Beta diversity

#### Bacterial community

PCA based on Aitchison distances revealed that fresh pulp samples clustered distinctly along PC1 (37.6% variance), separated from all fermented samples (Figure 4). At 48 h, zinc and amino acid treatments converged, while 24 h samples were more dispersed. Control samples at 24 h formed a separate cluster along PC2 (29.7%). PERMANOVA confirmed significant compositional differences between conditions (pseudo-F = 8.215, R^2^ = 0.691, *p* ≤ 0.01), with pairwise comparisons detecting significant differences between all condition pairs (*p* ≤ 0.05). Unweighted UniFrac PCoA confirmed the separation of fresh pulp from fermented samples (PERMANOVA: pseudo-F = 5.817, R^2^ = 0.613, *p* ≤ 0.01). Weighted UniFrac analysis further supported these patterns (PERMANOVA: pseudo-F = 19.069, R^2^ = 0.839, *p* ≤ 0.01), with the zinc treatment forming a distinct cluster.

**Figure 4.**
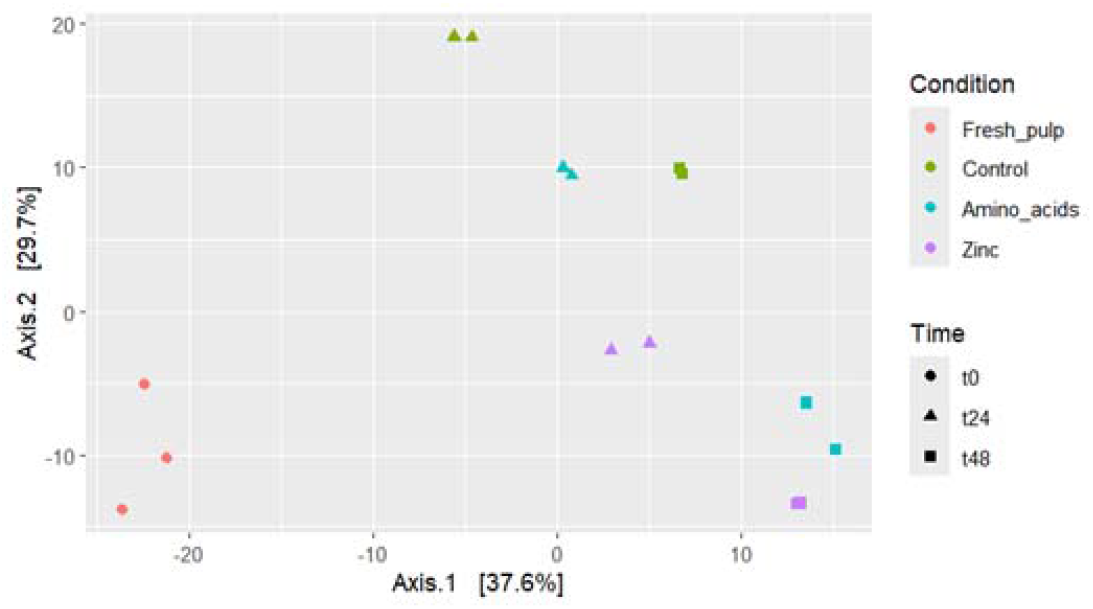
Principal component analysis (PCA) based on Euclidean distances from CLR-transformed bacterial ASV abundance data (Aitchison distance). Samples are colored by condition (Fresh pulp, Control, Amino acids, Zinc) and shaped by time point (t0, t24, t48).

#### Fungal community

PCA based on fungal Aitchison distances showed a distinct cluster for fresh pulp samples along PC1 (39.2%), while control samples at 24 h separated along PC2 from the supplemented treatments (Figure 5). ANOSIM confirmed significant differences among treatments (R = 0.359, *p* ≤ 0.05) and conditions (R = 0.571, *p* ≤ 0.01).

**Figure 5.**
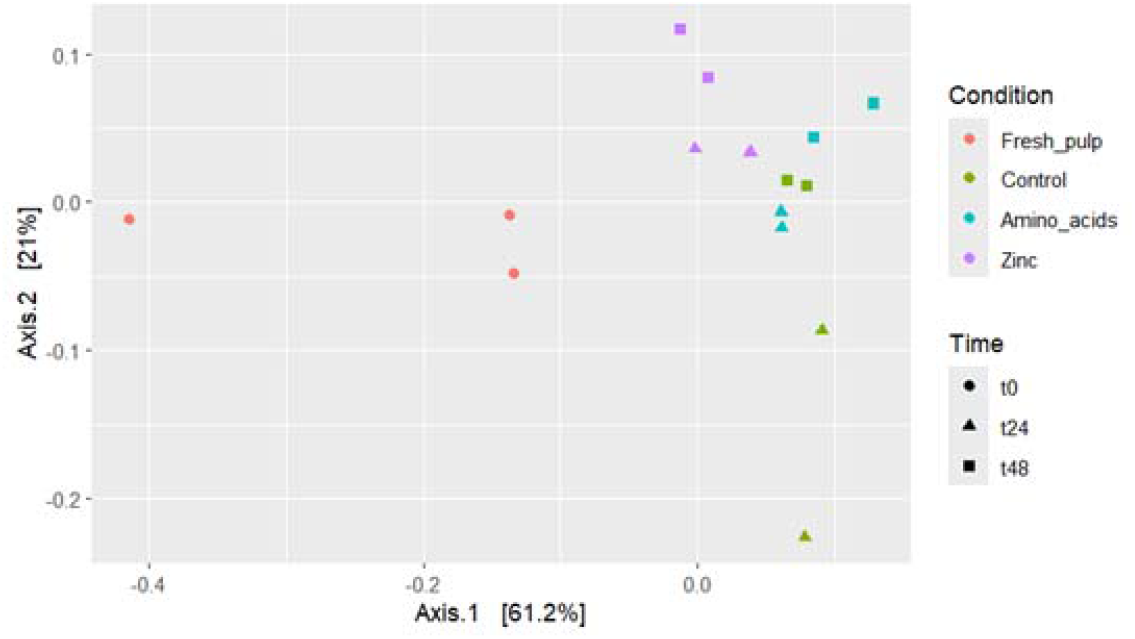
Principal component analysis (PCA) based on Euclidean distances from CLR-transformed fungal ASV abundance data (Aitchison distance). Samples are colored by condition and shaped by time point.

### 3.8. Functional prediction

PICRUSt2 analysis followed by DESeq2 testing revealed no significant differences in predicted KEGG pathway abundances between control and amino acid supplementation, or between control and zinc supplementation, at either 24 h or 48 h.

## 4. Discussion

To our knowledge, this is the first study to apply dual-kingdom metabarcoding to assess how nutritional supplementation affects the microbial ecosystem of cocoa fermentation within a geographically defined production territory. The results show that amino acid and zinc supplementation reshape microbial community composition and inter-taxon interactions without altering overall diversity, which points to targeted nutritional interventions as practical tools for modulating fermentation in cocoa-producing territories.

### 4.1. Baseline microbial diversity and territorial signatures

The bacterial and fungal diversity documented here is broadly consistent with previous findings from Costa Rican cocoa fermentations (Díaz-Muñoz et al., 2021; Van de Voorde et al., 2023). The dominance of *Weissella, Leuconostoc, Tatumella*, and *Acetobacter* among bacteria, and *Hanseniaspora, Pichia*, and *Thielaviopsis* among fungi, aligns with the established successional framework (De Vuyst and Weckx, 2016). The near-exclusive Proteobacteria dominance in fresh pulp (99.26%), driven largely by *Tatumella* (88.79%), is remarkable. This enterobacterium is associated with fresh plant tissues (Brady et al., 2013), and its rapid decline during fermentation suggests displacement by more competitive fermentative genera. The presence of *Cyphellophora* among fungi in the fresh pulp may represent a distinctive feature of this production territory. A recent study using genome-resolved metagenomics demonstrated that bacterial communities in cocoa fermentation are maintained by fermentation infrastructure (wooden boxes serving as microbial reservoirs), while fungal communities show stronger geographic signals (Gopaulchan et al., 2025). The territory-specific fungal assemblages observed here, including the unusual *Cyphellophora* abundance, are consistent with this model and suggest that the fungal fraction may serve as a marker of territorial identity.

### 4.2. Nutrient-specific modulation of community composition

Amino acid supplementation promoted a shift from Firmicutes (*Weissella, Leuconostoc*) toward Alphaproteobacteria (*Acetobacter*) at 48 h. This shift likely reflects enhanced AAB metabolic activity in the presence of amino acid-derived substrates: the Ehrlich pathway converts branched-chain amino acids to higher alcohols and acetate esters, and the intermediate products serve as substrates for *Acetobacter* oxidative metabolism (Hazelwood et al., 2008; Díaz-Muñoz and De Vuyst, 2022). In brewing, amino acid supplementation similarly accelerates transitions from LAB-to AAB-dominated phases (Bell and Henschke, 2005).

Zinc supplementation favored *Liquorilactobacillus* and *Lactiplantibacillus* among bacteria and *Wickerhamomyces* and Mucoromycota among fungi. The enrichment of Mucoromycota under zinc supplementation is noteworthy: members of this phylum possess diverse metalloenzymes and may benefit from enhanced zinc availability (Walker, 2004). *Wickerhamomyces* is recognized for its biocontrol potential and stress tolerance in fermented environments (Díaz-Muñoz et al., 2021), and its competitive advantage under zinc supplementation may relate to efficient zinc transport systems. The negative correlation between *Hanseniaspora* and *Wickerhamomyces* (−0.79) suggests competitive exclusion dynamics. Since these genera differ in their ester biosynthesis capacities (Crafack et al., 2013), zinc-induced shifts in their relative abundance could alter the volatile profile, broadening the aromatic complexity of fermented cocoa.

### 4.3. Community structure versus diversity

Beta diversity analyses consistently detected significant compositional differences between treatments (PERMANOVA R^2^ up to 0.839), yet alpha diversity indices showed no significant pairwise differences. This decoupling indicates that supplementation rebalances relative abundances and interspecific interactions without eliminating taxa. This is a desirable outcome for fermentation management. Interventions that reshape community function while preserving biodiversity are generally preferred, since species-rich communities tend to be more functionally resilient (Shade et al., 2012).

### 4.4. Ecological coherence of microbial interaction networks

SECOM analysis revealed ecologically coherent co-occurrence patterns. The positive correlation between *Gluconobacter* and *Acetobacter* (0.84) reflects their shared niche as obligate aerobic AAB during the later oxidative phase. The reciprocal negative correlations between *Tatumella* and multiple LAB/AAB genera suggest that the decline of this initial colonizer is a prerequisite for the establishment of the classical fermentation community. In the fungal community, the negative correlation between Ascomycota and Mucoromycota (−0.87) and between Saccharomycetes and Mucoromycetes (−0.89) indicates that zinc-induced Mucoromycota enrichment may come at the expense of the dominant Saccharomycetes. While Saccharomycetes are the primary ethanol and ester producers, Mucoromycota may contribute distinct enzymatic activities (e.g., pectinases, lipases) that enhance pulp degradation and precursor generation (Campos et al., 2025).

### 4.5. Limitations of functional prediction

The absence of significant differences in predicted KEGG pathway abundances may reflect inherent limitations of PICRUSt2, which relies on reference genome databases and assumes functional conservation among phylogenetically related taxa (Douglas et al., 2020). Functional redundancy, a hallmark of fermentation microbiomes, may mask compositional changes at the functional level (Gopaulchan et al., 2025). Additionally, sampling at 0, 24, and 48 h may be insufficient to capture transient functional shifts. Future studies integrating metatranscriptomics and targeted metabolomics (GC-MS volatile analysis) would enable more precise assessment of functional consequences.

### 4.6. Implications for quality differentiation in delimited territories

The capacity of nutritional supplementation to reshape microbial community structure without reducing diversity offers a practical tool for producing territories seeking to differentiate their cocoa through controlled fermentation. Within geographical indication frameworks, demonstrating that specific practices produce consistent, territory-linked microbial signatures and associated sensory attributes strengthens the scientific basis for quality certification (Hernandez and Granados, 2021). The distinct responses to amino acid versus zinc supplementation suggest that different strategies could be tailored to enhance specific aromatic profiles. For instance, amino acid supplementation could enhance fruity and banana-like notes via increased Ehrlich pathway flux, while *Wickerhamomyces*-mediated ester production under zinc supplementation could contribute more complex floral notes. Such tailored approaches would expand the sensory palette available to fine-flavor cocoa producers.

## 5. Conclusions

This study demonstrates that targeted supplementation with amino acids or zinc selectively modulates bacterial and fungal community structure during cocoa fermentation without altering overall microbial diversity. Amino acid supplementation favored *Acetobacter* and reduced Firmicutes dominance at 48 h, while zinc supplementation promoted Mucoromycota and *Wickerhamomyces* while sustaining *Liquorilactobacillus* abundance, indicating nutrient-specific ecological responses. These shifts reflect alterations in the balance between LAB and yeasts, the functional drivers of flavor precursor generation in fine cocoa. The structured co-occurrence networks revealed by SECOM underscore the ecological coherence of the fermentation microbiome and point to genera that may serve as biomarkers of supplementation effects. These results advance the understanding of microbiological diversity and differentiating factors in cocoa fermentation systems within delimited territories, and provide a foundation for evidence-based fermentation management. Future research should integrate metabolomic and metatranscriptomic analyses to link supplementation-induced community changes with the volatile compound profiles that define the sensory identity of territory-specific fine-flavor cocoa.

## Author Contributions

C.E.H. conceptualized the study, designed the experiments, and drafted the manuscript. A.M. contributed to data analysis, literature review, and manuscript revision. F.S.C. supervised field sampling, contributed to experimental design, performed bioinformatic and statistical analyses, and drafted the manuscript. A.A.L. coordinated field trials and sample logistics. All authors reviewed and approved the final manuscript.

